# Evidence for natural selection and barrier leakage in candidate loci underlying speciation in wood ants

**DOI:** 10.1101/500116

**Authors:** J Kulmuni, P Nouhaud, L Pluckrose, I Satokangas, K Dhaygude, RK Butlin

## Abstract

While speciation underlies novel biodiversity, it is poorly understood how natural selection shapes genomes during speciation. Selection is assumed to act against gene flow at barrier loci, promoting reproductive isolation and speciation. However, evidence for gene flow and selection is often indirect. Here we utilize haplodiploidy to identify candidate barrier loci in hybrids between two wood ant species and integrate survival analysis to directly measure if natural selection is acting at candidate barrier loci. We find multiple candidate barrier loci but surprisingly, proportion of them show leakage between samples collected ten years apart, natural selection favoring leakage in the latest sample. Barrier leakage and natural selection for introgressed alleles could be due to environment-dependent selection, emphasizing the need to consider temporal variation in natural selection in future speciation work. Integrating data on survival allows us to move beyond genome scans, demonstrating natural selection acting on hybrid genomes in real-time.

## Introduction

New species are formed when populations become reproductively isolated, i.e. they do not interbreed and exchange genetic material [1]. However, until complete reproductive isolation has evolved gene flow may still occur between the diverging lineages. This genetic exchange is usually localized in the genome, with some regions of the genome resisting gene flow better than others. Regions resistant to gene flow are thought to harbor “barrier loci” that can drive the divergence between lineages despite the homogenizing effect of gene flow [2]. Barrier loci could contribute to reproductive isolation by allowing differential adaptation between diverging lineages [2]. Alternatively, barrier loci could consist of genes incompatible between the diverging lineages (Dobzhansky-Muller incompatibilities [3,4], DMIs) driven by drift or divergent selection [5].

Several recent studies have searched for barrier loci using genome scans and identified genomic islands of differentiation using various measures of genetic differentiation [e.g. refs *6*–*8*]. The underlying assumption in these genome scans is that the most differentiated genomic regions between hybridizing lineages are those that are resistant to gene flow and hence drive reproductive isolation. Genetic differentiation however is an indirect measure of gene flow and several problems with genome scans have been discussed [e.g. refs *9, 10*]. The main problem lies in the fact that several other evolutionary forces (such as low recombination rate or drift) can bring about genomic regions of seemingly high differentiation between populations. Thus, we need more studies that test if natural selection is promoting speciation and halting gene flow at these genomic regions in natural populations.

Here we use ants as a model system to discover genomic regions of divergence (i.e. putative barrier loci) and to test for natural selection acting on them. In ants, males are haploid and born from unfertilized eggs from the mother, whereas females are diploid and arise via normal sexual reproduction. The haploid males thus allow the observation of natural selection acting on recessive alleles that are masked in female heterozygotes. Specifically, we use a naturally occurring hybrid population between two mound-building wood ant species, *Formica aquilonia* and *F. polyctena* from Southern Finland. These species have been estimated to diverge within the last 500, 000 years, probably at least partly in allopatry in different glacial refugia [10]. The hybrid population is well suited for unraveling the genetic basis of speciation: genetic incompatibilities segregate and persist across generations allowing to measure selection acting on these alleles in nature [11,12].

Previously, we have shown that haploid hybrid eggs (i.e. hybrid males) are laid each generation, but that males with specific introgressed alleles have low viability and die during development [12]. Yet diploid hybrid females with the same alleles persist in the population and seem to be favored over more parental-like individuals. This suggests recessive nuclear incompatibilities are in action in the haploid hybrid males, but these are masked in the diploid hybrid females where heterozygotes for introgressed alleles are favored [12]. Modeling work suggests that these opposite selective pressures between males and females can lead to purging of incompatibilities or their long-term maintenance depending on their initial frequencies, strength of selection and recombination [13]. The age of the hybrid population is unknown, but present-day individuals are not first-generation hybrids. Instead, the study population divides into two genetic lineages in a very similar way to the swordtail fish [14]; within the hybrid population one lineage is genetically more similar to one of the parental species, *Formica aquilonia* (previously referred to as W group), and the other is closer to the other parental species, *F. polyctena* (previously referred to as R group) [15]. Even though both lineages are of hybrid origin they show signs of reproductive isolation, as 94.5 % of successful matings are within a lineage [11,12] and we have never observed reproductive adult F1 individuals in nature. The *Formica* system differs from the genetic caste determination system observed e.g. in *Pogonomyrmex* harvester ants [16], because in *Formica* both queens and workers arise from the same mating events and show no genetic difference in contrast to *Pogonomyrmex*. Both lineages have previously lacked adult males with specific introgressed alleles (i.e. heterospecific alleles) or had them at low frequency due to negative selection, while the same alleles were favored in females as heterozygotes. This has resulted in strong allele frequency differences between sexes at specific marker loci, females having frequencies as high as 0.4 while males lack these alleles altogether [11]. We have shown that this antagonistic selection acting on introgressed alleles is stronger in the *F. polyctena*–like lineage, where over 90 % of male eggs were estimated to die [12]. Selection is weaker on the males of the *F. aquilonia*–like lineage, as is the evidence for selection favoring female heterozygotes.

Here we discover and annotate candidate barrier loci, i.e. the genomic regions that associate with hybrid male breakdown and thus act as barriers for gene flow between the two wood ant species. We then test if natural selection is acting on these candidate barrier regions in nature using SNP genotyping and survival analysis. This allows for testing the role of natural selection in genomic islands of divergence and reproductive isolation within a natural population, which has not been possible in most of the genomic studies of speciation.

## Results

### Multiple barrier loci throughout the genome are candidates for hybrid male breakdown

We used pooled genomic sequencing to discover the genomic regions associated with hybrid male breakdown (see Figure 1 for experimental design). We sequenced four samples, each consisting of 24 individuals: *F. polyctena*–like males, *F. polyctena*–like females (unmated queens), *F. aquilonia*–like males and *F. aquilonia* –like females (unmated queens). These individuals were classified into lineages based on six to seven diagnostic microsatellite alleles [11]. Due to haplodiploidy, this resulted in 24 and 48 chromosomes sampled for male and female pools, respectively. These pools represent population samples and came from two to four nests each, all of which belong to the same supercolony, where nests have hundreds of reproductive queens and relatedness approaching zero. Our previous [11,12] and current analyses (see Results) show there is no differentiation between nests within a lineage, justifying pooling of individuals from different nests. We sequenced each pool with 100 bp paired-end reads on its own lane in Illumina HiSeq2000 and made a *de novo* assembly from each pool. Subsequently, the assembly of *F. polyctena*–like males was used as a reference (genome size = 222.6 Mb, N50 = 1748 bp, 327,480 total scaffolds). Each pool was mapped to the reference assembly and SNPs were called after filtering. We retained 166,167 bi-allelic SNPs displaying coverage between 20 and 60 in each pool and minor allele count of four over all pools.

**Figure 1.**
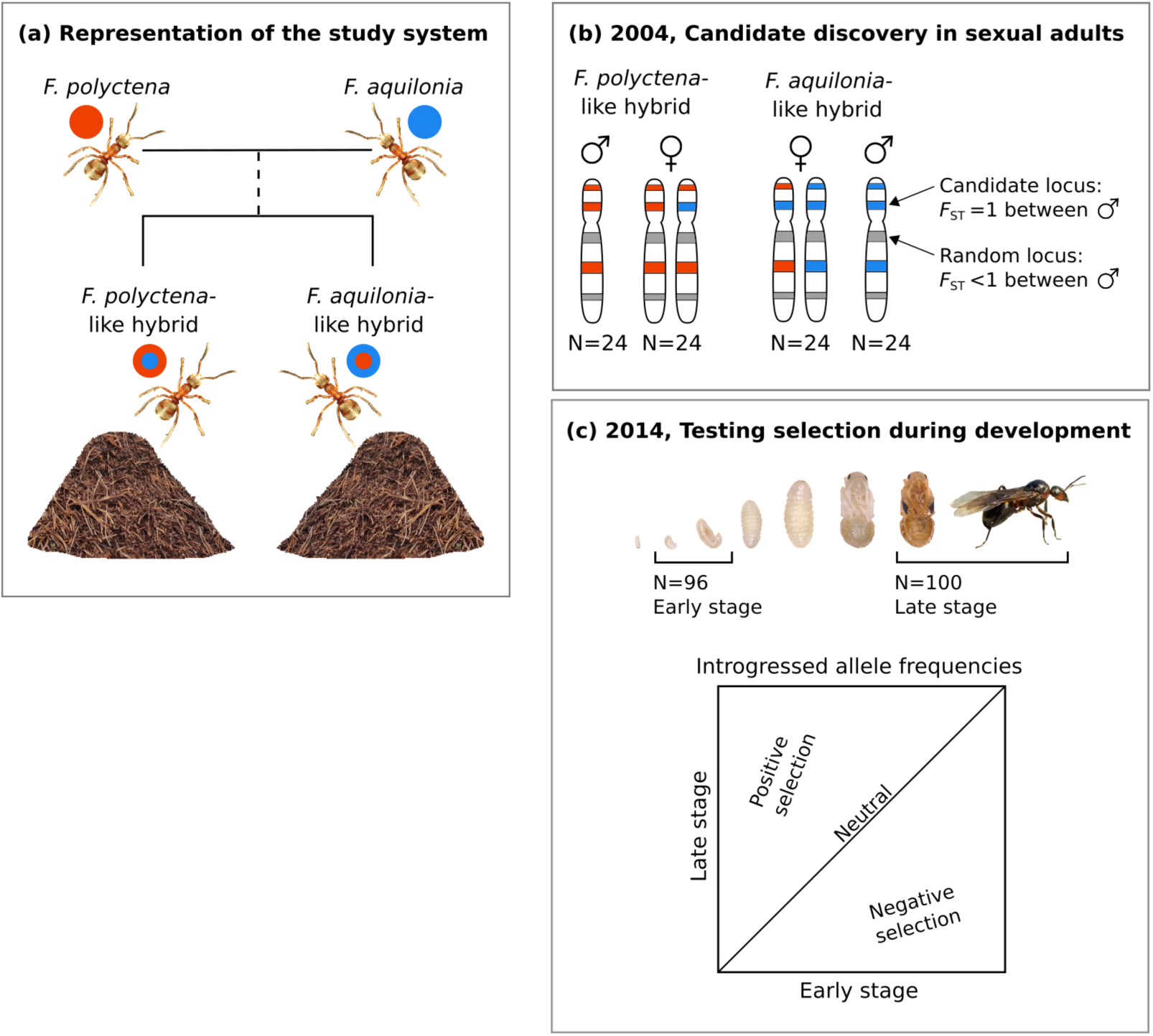
Overview of our approach. a) Study system, where two hybrid *F. aquilonia* × *F. polyctena* lineages coexist in different nests within the same population. b) Discovering candidate barrier regions with pooled genomic sequencing of adults, with a schematic representation of the male (haploid) and the female (diploid) genomes of the two lineages (simplified as a single chromosome). Red rectangles represent alleles originating from the *F. polyctena*-like lineage and blue represent alleles originating from *F. aquilonia-*like lineage. c) Testing natural selection at candidate barrier loci between larval (Early) and adult stages (Late).

We defined barrier loci as SNP positions where the male genomes from the *F. aquilonia*–like and *F. polyctena*–like lineages were fixed differently (i.e. *F*_ST_ = 1, see Supplementary Figure 1). This criterion is arbitrary in the sense that many loci showing significant differentiation (but not fixed) could be under selection too. However, loci fixed differently in males, despite shared hybrid history and variable in females are expected to experience the strongest antagonistic selection in the current population and offer the best opportunity to test for selection acting on barrier loci. We assumed that the allele fixed in *F. aquilonia*–like males represents the parental *F. aquilonia* allele and the allele fixed in *F. polyctena*–like males represents parental *F. polyctena*, from lineages that gave rise to the hybrid population.

The mean *F*_ST_ estimate between the *F. aquilonia*–like and *F. polyctena*–like lineages was 0.14 (s.e. = 4.66×10^−4^) between the males, and 0.07 (s.e. = 2.27×10^−4^) between the female pools. We found 711 SNPs (0.43 % of the total) alternatively fixed between males of the *F. aquilonia*–like and *F. polyctena*–like lineages, but no fixed differences between the female pools. As expected under antagonistic selection, SNPs fixed differently in males were polymorphic and had mean *F*_ST_ of 0.34 (s.e. = 5.83×10^−3^, Supplementary Figure 1) in the females. The alternatively fixed SNPs between the male pools represent genomic regions of high differentiation between the two male lineages despite their shared hybrid history, and thus mark candidate barrier loci that are predicted to contribute to reproductive isolation and speciation. These 711 SNPs fell into 610 assembled contigs (cumulative size: 2.09 Mb, 0.94% of the genome), whose lengths vary between 515 – 23,815 bp (mean = 3430 bp). Seventy-eight contigs had more than two barrier SNPs (maximal number of barrier SNPs on a single contig was 5).

We were interested in how these candidate barrier loci locate in the genome, because this can provide information about the origin of reproductive isolation and play a role in how efficient the barrier loci are in maintaining divergence despite hybridization and gene flow. If candidate barrier loci are co-localized in the genome their origin and maintenance could be explained by selection on a few genomic regions that could, for example, harbor structural re-arrangements. Alternatively, candidate barrier loci scattered across the genome could effectively reduce gene flow over much of the genome. We used the genome of the closely-related *Formica exsecta* [17] (277 Mb, 14,617 scaffolds, N50 = 997,654), which is better assembled compared to the poolseq genome assembly, to map the location of candidate barrier loci. *F. exsecta* is estimated to have diverged from the parental species of our hybrid population 5 Mya [18]. We used the 610 contigs containing candidate barrier SNPs as queries and searched the *F. exsecta* genome assembly using BlastN. If our query scaffold was larger than 5, 000 bp we used +/-2, 500 bp surrounding the candidate barrier SNP as query. Our candidate barrier loci map to a total of 134 *F. exsecta* contigs, with a cumulative coverage of 56.2 % of the *F. exsecta* genome, both of which are significantly less than for a similar number of random SNPs in our data set (Supplementary Figure 2). Haploid chromosome number varies between 26 and 28 in *Formica* [19]. Assuming evenly-sized chromosomes our candidate barrier loci are likely to be situated in over 10 chromosomes.

**Figure 2.**
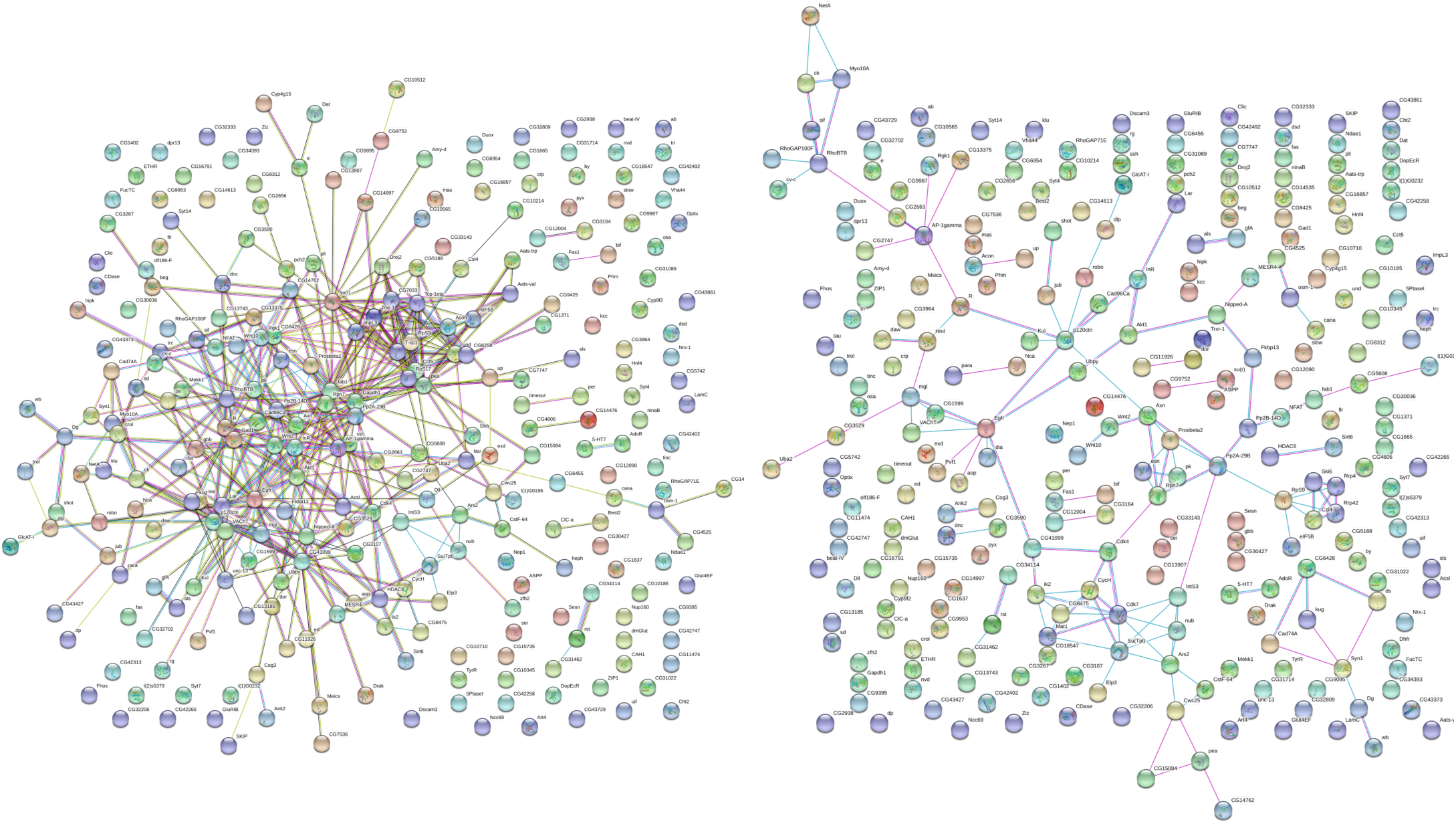
STRING interaction network. Each circle represents one candidate gene and lines connecting them indicate support for interaction between the genes. Majority of interactions fall into a single network. Line color indicates following: blue = data from curated databases, pink = experimentally determined interactions, black = co-expression. Analysis is based on *D. melanogaster* homologs. Left panel: all interactions, and right panel: experimentally determined interactions or interactions obtained from curated databases.

### Candidate barrier loci are predicted to interact at the protein level

Next, we were interested in the possible functions of our candidate barrier loci, as they can be informative about the putative selection pressures and mechanisms of hybrid male breakdown. We annotated each of the 610 contigs containing candidate barrier SNPs, first extracting the transcripts of the genes located in those regions from the closely related *F. exsecta*. Out of 571 candidate barrier regions (see Material and Methods; each region was defined as 5, 000 bp around the candidate barrier SNP) that we were able to anchor in *F. exsecta* (all blastn *e*-values < 10^−141^), we recovered 590 unique genes. Sixty-three of the candidate genomic regions did not contain any annotated gene. The mean number of genes per candidate genomic region was 1.4, and the maximum was six. For each gene, its *F. exsecta* transcript sequence was recovered and blasted (blastx) against *Drosophila melanogaster* r6.15 proteins recovering 317 genes, whose homologs reside in candidate barrier regions (Supplementary Table 1).

The genetic model for hybrid breakdown suggests inviability and sterility are caused by dysfunction and negative epistatic interactions between diverged genes from the parental species. One possible mechanism that can result in epistasis are molecular interactions. We thus asked if there was any evidence for molecular interactions in the form of protein-protein interactions between the gene products in candidate barrier regions. For this we used the recovered *D. melanogaster* proteins and the STRING database [20], which contains information on known and predicted protein-protein interactions. We found that candidate barrier regions harbor proteins that show significantly more interactions with each other than a random set of proteins drawn from the *D. melanogaster* genome (PPI enrichment *p*-value = 9.65×10^−13^, Figure 2). Out of 268 genes annotated and having information in the STRING database, 167 genes fall into a single interaction network, with evidence for protein-protein interactions in *D. melanogaster* or other species. Seventy-four of the network genes had evidence for interactions which were experimentally determined or obtained from curated database (PPI enrichment *p*-value = 0.03). However, randomly drawing 711 SNPs from our pooled sequencing data 1000 times showed that similar enrichment in protein-protein interactions (i.e. network) could be retrieved for a random set of SNPs as well. This suggests either a bias towards functional and interacting sets of genes in our total SNP dataset or a bias in our method towards interacting sets of genes arising due to conservation (only genes conserved between ants and *D. melanogaster* can be included in the analysis). However, this does not mean the interactions which were documented would be unreliable. In summary, our results suggest interactions among the genes in candidate barrier regions, but these interactions are not significantly more abundant than for random sets of SNPs in our data.

Next, we were interested in the functions of genes within candidate barrier regions. For this we used the *D. melanogaster* gene IDs. The top three biological processes characterizing our candidate barrier genes are “developmental process” (*p* = 3.5×10^−18^), “system development” (*p* = 3.5×10^−18^), and “single-organism developmental process” (*p* = 4.03×10^−18^). Similarly, the top three molecular functions are “ion binding” (*p* = 8.55×10^−6^), “protein binding” (*p* = 3.67×10^−5^) and “calcium ion binding” (*p* = 8.74×10^−5^). These biological processes and molecular functions were highly significantly enriched when compared to the genome average of *Drosophila melanogaster*. However, they were not significantly enriched if compared to a random set of SNPs drawn from our data, as all of them are found in over 5 % of our simulations.

### Testing for signs of selection at candidate barrier loci in a natural population

We tested for selection during development by genotyping altogether 96 individuals at early (larva) and 100 individuals at late (adult or late stage pupa) developmental stages (Supplementary Table 2), with 180 random and 183 candidate barrier SNPs (Figure 1). Out of these, 163 candidate barrier SNPs and 137 random SNPs were successfully genotyped. The 137 random SNPs were located on 136 contigs (cumulative size: 484 kb, 0.22% of the genome), whereas the 163 candidate barrier SNPs located on 159 contigs within our assembly (cumulative size: 523 kb, 0.23% of the genome). Individuals used in genotyping were collected in Spring 2014 from the same hybrid population as the one sampled for the pooled sequencing analysis ten years earlier.

The SNP genotyping confirmed that within the hybrid population, *F. polyctena*–like and *F. aquilonia*–like lineages are genetically distinct from one another (Supplementary Figure 3). We found two small female larvae, which were genetically intermediate and had unique combinations of putative parental alleles from the two lineages suggesting they were early-generation hybrids between the lineages (Supplementary Figure 3). The *F. polyctena*–like individuals, and especially the males, were previously shown to experience strongest selection. Unfortunately, we were able to sample only 8 males from this lineage, five of which turned out to be diploid, thus *F. polyctena*–like males were excluded from the analysis. The following analyses were performed for *F. polyctena*-like females, *F. aquilonia*-like males and *F. aquilonia*-like females.

**Figure 3.**
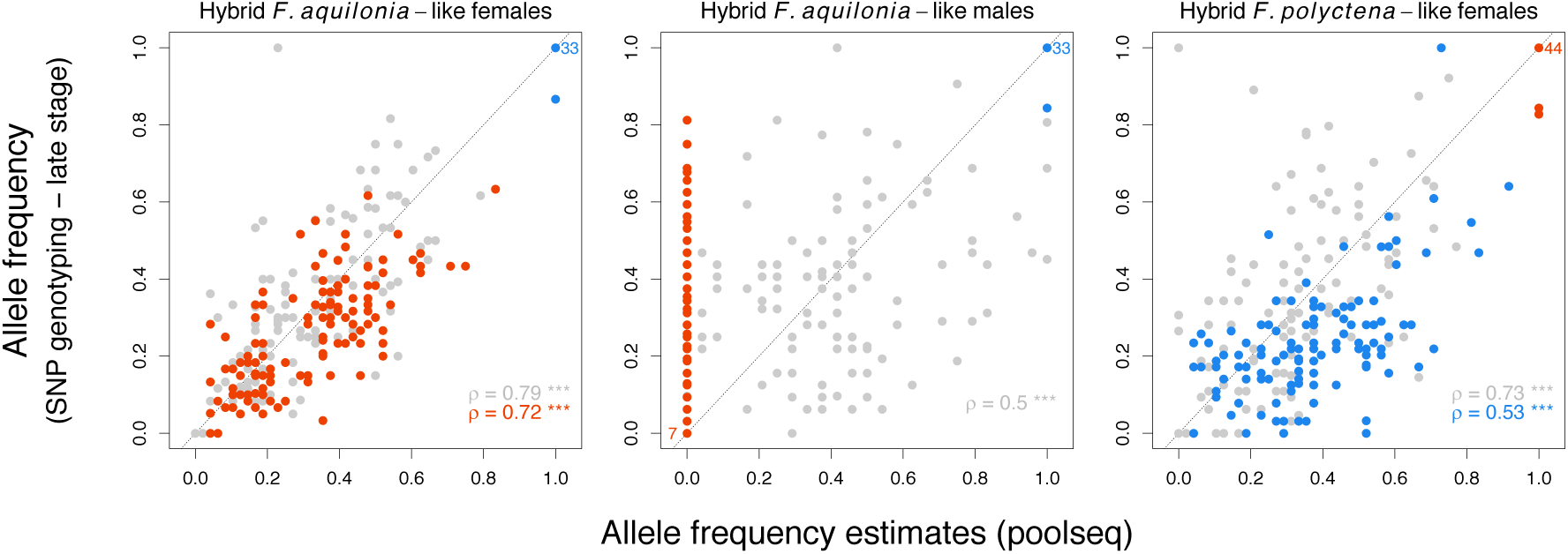
Allele frequencies correlate between pooled sequencing (adults sampled in 2004) and SNP genotyping in late stage (adults sampled in 2014). Putative barrier loci are colored according to their polarization (red: allele introgressed from *F. polyctena*-like to *F. aquilonia-*like, blue: allele introgressed from *F. aquilonia*-like to *F. polyctena*-like). Control SNPs (in grey) are polarized by the minor allele in the poolseq data. Linear regressions are indicated for each class of SNPs according to their colors, along with their Spearman’s correlation coefficients and their significance (***: *p* < 0.001).

First, we asked if adult allele frequencies inferred from the pooled sequencing correlated with adult allele frequencies estimated from the SNP genotyping data, as the two data sets have been collected ten years apart and allele frequencies were estimated in different ways. Over all the genotyped SNPs, allele frequencies between the two data sets correlated well in *F. polyctena*–like females (Spearman’s correlation, ρ = 0.87, CI_95_ = [0.82, 0.90], *p* < 0.001) and in *F. aquilonia*–like females (ρ = 0.91, CI_95_ = [0.88, 0.94], *p* < 0.001) (Figure 3). The correlation was lower for the males with a Spearman’s correlation coefficient of 0.71 (CI_95_ = [0.64, 0.76], *p* < 0.001) between adult allele frequencies estimated from pooled sequencing and genotyping (Figure 3). The lower correlation in males compared to females can be explained, at least in part, by the lower number of sampled chromosomes for (haploid) males compared to (diploid) females creating a larger error in the estimated allele frequencies. Overall, correlations between allele frequency estimates are in line with other studies focusing on the accuracy of allele frequency estimates derived from poolseq [21] and suggest pooled sequencing and SNP genotyping gave similar results.

### Candidate barrier SNPs show significant heterozygote excess in females

Next, we tested if the barrier loci showed significant allele or genotype frequency changes during development from larva to adult compared to the random loci. This would demonstrate natural selection acting on the candidate barrier loci in nature. No significant allele frequency change was observed between larval and adult stage at candidate barrier loci compared to random loci in females of *F. aquilonia*– or *F. polyctena*–like lineage (Figure 4, Figure 5).

**Figure 4.**
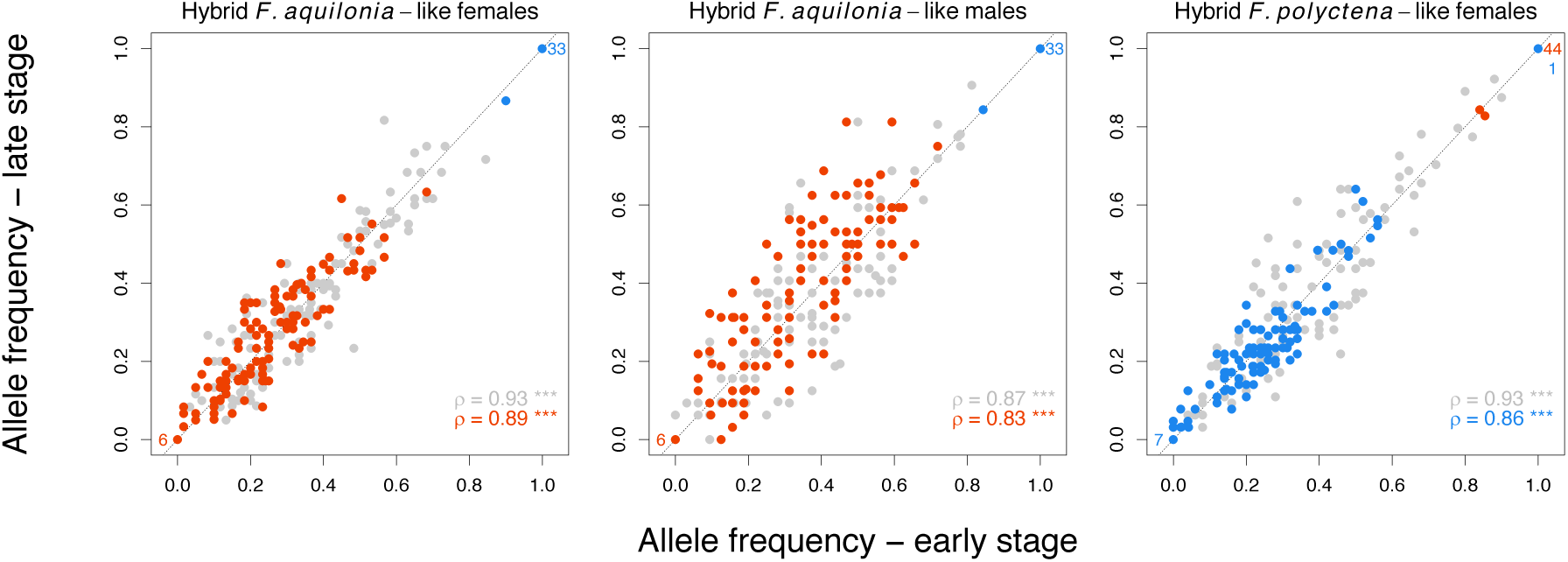
Allele frequency correlation between early (larva) and late (adult) developmental stages in random (grey) and candidate barrier loci (red: allele introgressed from *F. polyctena*-like to *F. aquilonia*-like, blue: allele introgressed from *F. aquilonia*-like to *F. polyctena*-like) in year 2014. No strong allele frequency change was observed from larva to adult in females, but frequencies in males were significantly different between larva and adult (see results). Numbers in bottom left and upper right parts indicate the number of candidate barrier SNPs fixed. Linear regressions are indicated for each class of SNPs according to their colors, along with their Spearman’s correlation coefficients and their significance (***: *p* < 0.001).

**Figure 5.**
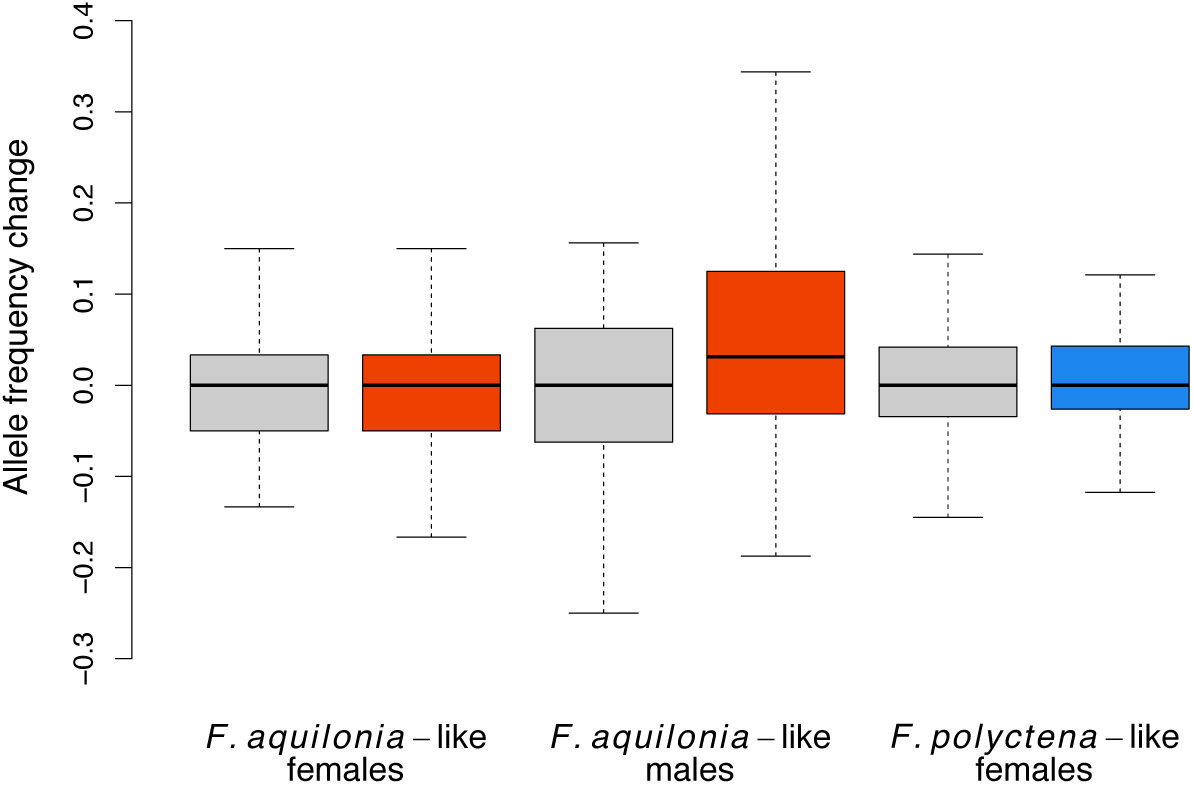
Mean allele frequency change between early (larva) and late (adult) developmental stages in random (grey) and candidate barrier loci (red or blue) in year 2014. Allele frequencies at barrier loci are polarized according to the introgressed allele. Candidate barrier SNPs have significantly stronger allele frequency change compared to random SNPs in *F. aquilonia*–like males (*p* = 0.003, see methods for details on the model).

Our previous microsatellite-based study showed that introgressed alleles were favored in females when heterozygous. This selection acted throughout the female lifetime. Thus, we searched for signs of similar selection testing for increase in heterozygote excess between early and late stages using *F*_IS_ estimates. Indeed, there was a tendency for more negative mean *F*_IS_ values (i.e. greater excess of heterozygosity) in candidate barrier loci at the adult stage (*F. aquilonia*–like *F*_IS_ = –0.116, *F. polyctena*– *F*_IS_ = – 0.2) compared to the larval stage (*F. aquilonia*–like *F*_IS_ = –0.085, *F. polyctena*– *F*_IS_ = –0.179) in both female lineages (Figure 6), but this tendency was not statistically significant in either lineage when compared to random loci (*F. aquilonia*–like *p* = 0.295, *F. polyctena*–like *p* = 0.643). However, candidate barrier loci had significantly greater heterozygote excess compared to random loci already at larval stage (GLMM, *z*-value = –4.404, *p* < 0.001) and this pattern remained at adult stage as well (GLMM, *z*-value= –7.133, *p* < 0.001) in *F. polyctena*–like females. To account for possible effects of linkage disequilibrium, the SNP data set was pruned by omitting SNPs closer than 50 kb, 100 kb or 500 kb according to their BLAST hits in the *F. exsecta* assembly. Pruning did not affect the above results (all *p*-values reported above from GLMMs remained significant for all pruning intervals; Supplementary Figure 8, panels A-D). Greater excess heterozygosity in candidate barrier SNPs compared to random SNPs can result from either (i) strong allele frequency differences between parental genotypes or (ii) selection for heterozygosity that has already acted at candidate barrier loci before the larval stage we sampled (i.e. between egg and larval stage).

**Figure 6.**
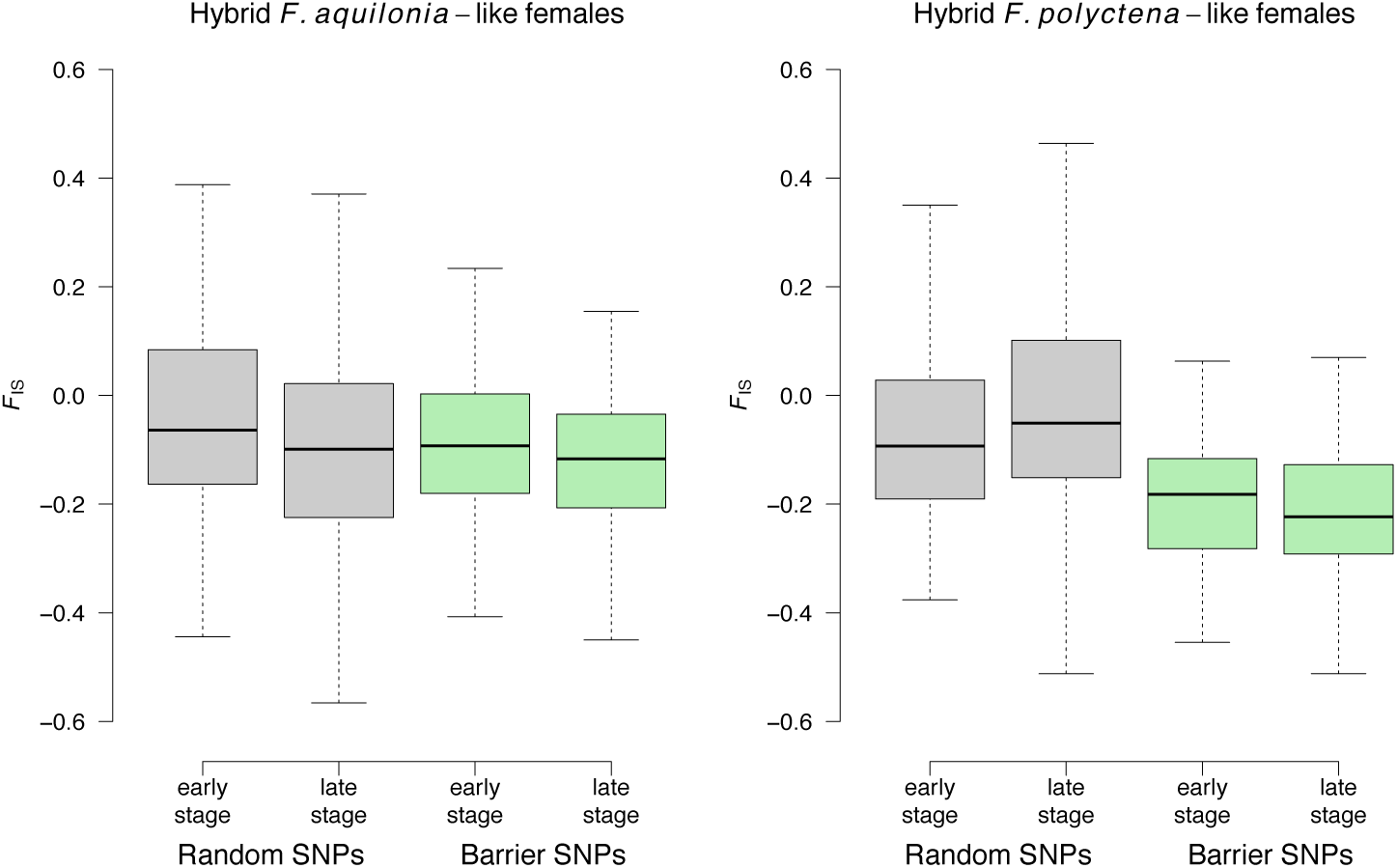
Mean F_IS_ estimates in early (larva) and late (adult) developmental stages in random (grey) and candidate barrier loci (green) in year 2014.

### Evidence for selection in candidate barrier SNPs in F. aquilonia–like males

Then, we tested for allele frequency changes in *F. aquilonia*–like males, expecting the introgressed alleles to decrease in frequency during development as shown in our previous microsatellite based study [12]. Allele frequencies were significantly different between larval and adult developmental stages in candidate barrier SNPs compared to the random SNPs (glmer, *z*-value= –2.98, p = 0.00286), the candidate barrier SNPs showing more frequency change during development (Figure 5). However, the frequencies of the introgressed alleles increased from larval to adult developmental stages in *F. aquilonia*–like males (Figure 4), a pattern opposite to our expectations. Significant increase of introgressed alleles at candidate barrier loci in contrast to alleles at random loci suggests selection favored the introgressed alleles in *F. aquilonia*–like males during development. Again, this result remained significant after pruning the SNP data (Supplementary Figure 8, panel E). Unfortunately, we were unable to test selection acting at candidate barrier loci in *F. polyctena*–like males, the lineage where we had the strongest expectation of negative selection, because too few *F. polyctena*–like males were sampled from the population. The low occurrence of these males is consistent with their estimated high mortality [12].

### Barrier leakage and significant variation in the strength of barriers through time

Introgressed (i.e., *F. polyctena*–like) alleles were absent from adult *F. aquilonia*–like males at 163 candidate barrier loci in year 2004 (allele frequency of zero in the poolseq data). However, in year 2014 significant proportion of these loci harbored alleles introgressed from the *F. polyctena*–like lineage (123 loci, i.e. 75.5%) to *F. aquilonia*–like males. The mean frequency of introgressed alleles in these males was 0.36, the most common introgressed allele being at a frequency of 0.81. Sampling different sub-populations within a lineage at different years is unlikely to drive increase of introgressed allele frequencies between 2004 and 2014 in our samples of *F. aquilonia*–like males: We observed no population substructure within the *F. aquilonia*–like lineage in 2005 or 2014 (Supplementary Figures 3 and 4). Increase of introgressed alleles in males is not likely to be caused by errors in male poolseq-based allele frequency estimates either. All but one of the candidate barrier SNPs (33 out of 34 SNPs) that were initially fixed in both adult males and females of the *F. aquilonia*–like lineage in the pooled sequencing data (year 2004) were still fixed in the genotyping data (year 2014), showing consistency across data sets. If differences between allele frequency estimates from 2004 and 2014 were caused by errors in poolseq allele frequency estimates, this should affect all SNP loci equally. However, this is not the case, as only candidate barrier SNPs that were polymorphic in females of the *F. aquilonia*–like lineage have now “leaked” into *F. aquilonia*–like males. Our results thus suggest that in year 2014 a proportion of the barrier loci are “leaking” in *F. aquilonia*–like males. At these loci introgressed alleles were on average selected for in males as evidenced by significant frequency increase between larval and adult stages (see above), a pattern opposite to our previous observations [11,12].

Using microsatellite genotyping, we further verified that increasing frequencies of introgressed alleles at candidate barrier SNPs in *F. aquilonia*–like males did not represent technical artifacts. Specifically, we genotyped the individuals collected in 2014 and used in SNP genotyping with nine previously used microsatellite markers [5,12] and compared them to samples collected in 2004, 2008 and 2011 (Supplementary Table 3). Microsatellite allele frequencies from 2014 paralleled the pattern observed in candidate barrier SNPs in the same year. Indeed, *F. aquilonia*–like males in 2014 harbor microsatellite alleles introgressed from the *F. polyctena*–like lineage that were missing from them in 2004 (Table 2). Specifically, the frequencies of two microsatellite alleles (Fy3_190_ and Fy15_222_) introgressed from *F. polyctena* to *F. aquilonia*–like males have fluctuated over the years from 0 in 2004 to 0.28 in 2014 for Fy15_222_ and from 0 in 2004 to 0.45 in 2014 for Fy3_190_ (Supplementary table 3, Figure 7A). No leakage of alleles introgressed from the *F. aquilonia*-like lineage to the *F. polyctena*–like males was observed, the five introgressed microsatellite alleles remaining at a null frequency in adult *F. polyctena*–like males in 2004 (N=35), 2008 (N=41) and 2011 (N=23, Supplementary table 3).

**Figure 7.**
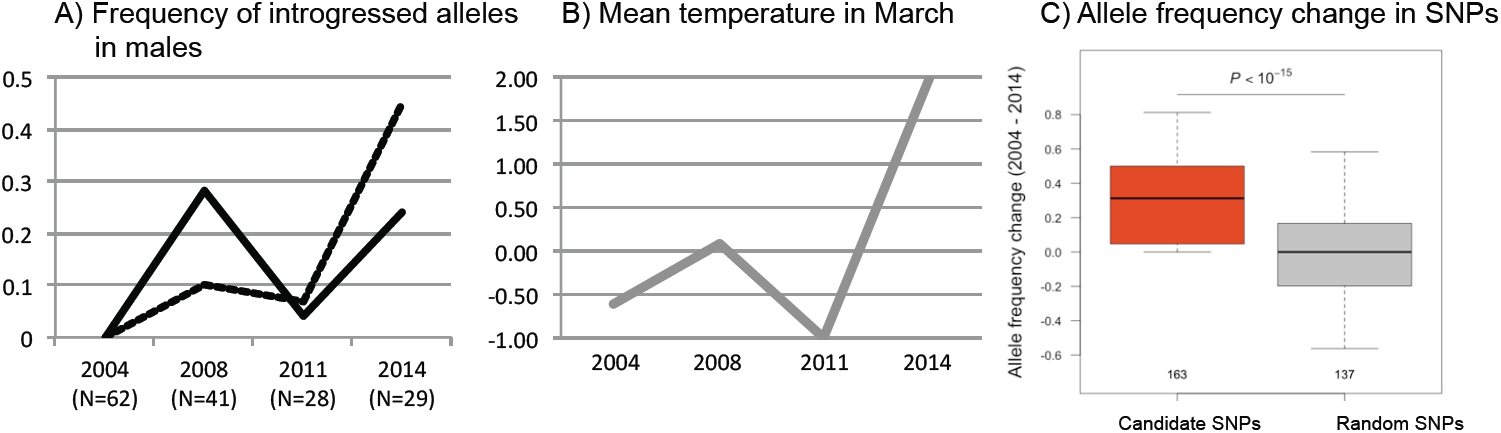
Frequency of introgressed alleles in adult *F. aquilonia*–like males appears to co-vary with yearly spring temperature. A) Frequency of alleles introgressed from *F. polyctena* to *F. aquilonia*–like males at two microsatellite loci (dashed FY13_190, straight FY15_222) in years 2004, 2008, 2011, 2014. B) Mean temperature (°C) in March during male development each year (Finnish Meteorological Institute, data from the nearest measurement point, Helsinki). C) Introgressed alleles at candidate SNPs (N=163) increase significantly more in frequency than alleles at random loci (N=137) between 2004 and 2014.

## Discussion

Several recent genome scan studies have used next-generation sequencing to map genomic regions putatively under divergent natural selection and underlying reproductive isolation (reviewed in [2]). Approaches can often be based solely on comparing genomes, because observing natural selection, collecting data on fitness or molecular characterization of loci underlying reproductive isolation is challenging.

However, several suggestions have been made to move beyond genome scans with an integrative approach [2,22]. To bridge the gap between genotype and fitness in genome scan studies we first took advantage of male haploidy in ants and discovered genomic regions of high divergence (i.e. candidate barrier regions). We then tested if natural selection is acting on these genomic regions in a natural population using a SNP panel and survival analysis, expecting haploid males to show the strongest signals of selection against gene flow at barrier loci. However, instead we document signature of barrier leakage at the candidate loci, with significant increase in the frequency of introgressed alleles in *F. aquilonia*–like males during development. Below we discuss reasons for the signatures of barrier leakage including environment-dependent incompatibilities, formation of compatible combinations of parental alleles and recombination breaking association between selected and hitchhiking loci. The ant system is dynamic and allows further investigations into how selection acts on hybrid genomes in cases of speciation with gene flow.

### Considering barrier loci that evolve in a network

Hybrid breakdown is expected to result from deleterious epistatic interactions (i.e. intrinsic incompatibilities) between diverged genes of the parental species [3,4]. Relatively few studies have investigated the effect of incompatibilities in networks or incompatibility loci with more than one interaction (in comparison to two-locus DMIs) for speciation, but see [5,23–25]. This is in stark contrast with our understanding of molecular biology: genes and their products interact with several other genes, are part of gene interaction networks, and are regulated by transcription factors. We found that candidate barrier and hitchhiking loci likely cover a large proportion of the genome in two *Formica* ant species, which diverged within the last 500 000 years [10]. We also found support for multiple protein-protein interactions between genes located among the candidate barrier regions. Majority of genes in these regions were predicted to form a single protein-protein interaction network, with interactions for homologous proteins documented or predicted in other species. However, our bootstrap analysis shows that the network found among the candidate barrier loci was not significantly different from the genomic background: any polymorphisms we were able to call within our pooled sequencing data was likely to give significant enrichment of protein-protein interactions using our analysis pipeline. Our annotation is based on *D. melanogaster* homologs, which makes our pipeline biased towards conserved genes. If conserved genes are also more likely to have evidence for protein-protein interactions (as suggested by some studies [26]), that could lead to enrichment of interactions among both candidate barrier loci and random sets of SNPs with our data and pipeline. Our results highlight potential biases when extrapolating data from model organisms into non-model systems. Bootstrapping analyses, such as presented here, can reveal that candidate genes are not any different from genomic background of the focal species. However, this does not mean that the interactions we found would be unimportant. Protein-protein interactions could still create epistatic interactions among the barrier loci. Our results are consistent with the idea of multiple genes underlying hybrid breakdown that interact in a network instead of multiple two-locus interactions. Functional and population studies in *Formica* are needed to test if mismatching combinations of parental alleles at network loci lead to hybrid inviability, as predicted if protein-protein interactions cause negative epistasis [3,4].

Annotation of candidate barrier regions suggests genes in these regions play a role in development, with 133 genes out of 268 investigated having the GO term “developmental process”. These results are consistent with our previous findings where we show hybrid male breakdown during development in the *F. polyctena*–like lineage [12]. Furthermore, developmental problems are a common cause of hybrid breakdown [27,28]. Our analysis pipeline is likely to produce a bias towards conserved genes because it is based on *D. melanogaster* homologs, and developmental genes are more likely to be conserved at the sequence level [29]. Thus, it is likely that additional gene functions could be found if we were able to annotate the function of all open reading frames within candidate barrier regions. Interestingly, our candidate barrier loci included homologs of two major barrier loci in *Heliconius*: a member of the *Wnt* gene family and *optix*, which are known to act as barriers to gene flow and are responsible for variation in wing patterning between *Heliconius* races [6].

But why would a developmental gene network have diverged between the two *Formica* species parental to the hybrids? Divergence could have been driven by drift or differential adaptation between the parental species. The parental species have likely diverged in allopatry and may now be adapted into different climatic conditions, *F. aquilonia* being distributed in Northern Eurasia and higher altitudes of Central Europe, while *F. polyctena* is likely adapted to warmer climate and occurs in Central Europe and Southern parts of Fennoscandia and Russia [30]. While the species have adapted into different climatic regimes it is plausible that their developmental programs have adapted and diverged too. However, developmental programs can diverge also at least partly by drift. Developmental system drift is a common phenomenon [31] where homologous characters develop via different regulation of genes or involve completely different sets of genes.

### Signs of selection at candidate barrier loci and barrier leakage

Previously we have shown antagonistic selection on ant hybrids, where introgressed alleles were favored in female heterozygotes, but selected against in haploid males, with strongest evidence of this in the *F. polyctena*–like lineage [11,12]. Our current dataset shows a trend of increasing heterozygosity excess at candidate barrier loci during female development, but this is not statistically significant after removal of two outlier individuals likely to be early generation hybrids. However, we find a significant excess of heterozygosity at the candidate barrier loci compared to the random loci. This excess heterozygosity is likely to be caused by selection either in the current generation or earlier in population history. First, strong allele frequency differences between mothers and fathers would result in high heterozygosity in diploid females without the action of selection in the sampled generation. However, in this scenario initial differentiation between sexes at candidate barrier loci was likely caused by selection acting in earlier generations (see below). Second, selection that favors heterozygotes may have already acted on the candidate loci before the larval stage we sampled. Our previous study documented selection during development using eggs and few days old larvae and compared them to adults, whereas here we use 1-2 week old larvae and adults. Thus, in the current study we are likely to miss any signal of selection acting earlier in development.

We assumed the candidate barrier loci to represent recessive incompatibilities and loci linked to them, thus expecting to find evidence of negative selection in the haploid males [11,12]. However, we find the opposite with introgressed alleles increasing in frequency at the candidate barrier loci in the *F. aquilonia*–like males during development. This is consistent with natural selection favoring introgression in these males. The magnitude of change is comparable to that observed within a generation in other systems, e.g. in stick insects for the color pattern locus [32].

Selection for introgression within a generation in males may be connected to patterns we observed between years: A proportion of the barrier loci showed barrier leakage resulting in increase of introgressed alleles in males sampled 10 years apart. These rising frequencies of introgressed alleles in *F. aquilonia*–like haploid males between years can be explained by at least three hypotheses, which are not mutually exclusive. First, the barriers to gene flow can be gradually breaking down in *F. aquilonia*–like males, which could explain both within- and between-year patterns. Barrier breakage could be due to recombination and subsequent formation of compatible combinations of barrier loci in males. Barriers for gene flow could also break down due to relaxed selection. Under this model the haplodiploid systems (and X-chromosomes) can show fluctuating allele frequencies at candidate barrier loci, comparable to those observed in our data, if sexes had different allele frequencies to begin with. However, gradual breakdown of barriers for gene flow can only explain changes observed between years and not selection for introgressed alleles within a generation in males. Also, allele frequency differences between sexes are not expected to be maintained without selection over multiple generations. Second, a proportion of marker loci may have been hitchhiking due to close genomic location with barrier loci and so missing from males in 2004, but these hitchhikers are now becoming dissociated from the barrier loci via recombination and consequently found in males in 2014. Again, this hypothesis cannot explain the significant increase of introgressed alleles within a generation in males. A third possibility is that the incompatibilities could be environment-dependent, which could explain both within-generation and between-years effect. Environment-dependence means that the negative effects of the incompatibilities would be expressed only in certain environments. This situation is comparable to genotype × genotype × environment interaction, where fitness effects of a genetic variant are dependent on the genomic background it is found in, as well as the external environment. Previous empirical evidence for environment-dependent incompatibilities comes e.g. from yeast [33] and *Drosophila* [34]. Current results for ants are most consistent with environment-dependent incompatibilities, but new genome-wide data needs to be collected over multiple years and generations in order to rule out other hypotheses.

### Can environment drive barrier leakage?

Environment dependent incompatibilities have been largely neglected in the speciation context: DMIs are assumed to be deleterious, no matter the environment. Yet, we know cases where barriers for gene flow hold in one environment but not in the other [14,33,34]. The fact that we see fluctuations rather than directional changes in the frequency of introgressed alleles at microsatellite loci (years 2004, 2008, 2011 and 2014) in the ant system is consistent with the idea of environment-dependent incompatibilities. What could be the selective agent? Temperature is one possible environmental variable that could affect the expression of incompatibilities due to two reasons. First, temperature differs between the current environments of the parental species, *F. aquilonia* being more Northern and *F. polyctena* being more Southern species. Second, temperature can show fluctuation between and within years. Indeed, the frequency of introgressed alleles in *F. aquilonia*–like males appears to co-vary with temperature during developmental season, but due to the limited time-series this correlation cannot be tested statistically (Figure 7). Further studies are needed to show if temperature is a key selective pressure that creates variation in the survival of ant hybrids. These analyses allow testing if incompatibilities are environment-dependent, an aspect that has been to a large extent ignored in speciation studies.

### Conclusions

Here we bridged the gap between genome scan studies and fitness by mapping candidate barrier loci between two recently diverged wood ant species and tested for natural selection acting on candidate barrier regions. We do document selection at candidate barrier loci, but it is acting on opposite direction than expected, favoring introgression in males. This barrier leakage may depend on the environment and fluctuate between years. These results highlight the dynamic nature of the ant system that allows investigations into genomic and molecular consequences of hybridization, areas where many questions still remain to be answered.

## Materials and Methods

### Experimental design

Our first aim was to use a genome scan approach to discover candidate barrier regions, i.e. genomic regions that drive divergence between two wood ant species despite of hybridization and gene flow. For this we used pooled whole genome sequencing. Our second aim was to directly measure if natural selection is promoting divergence and so speciation in the current-day hybrid population at the candidate barrier regions using survival analysis and SNP genotyping.

### Discovery of candidate barrier regions with pooled genomic sequencing

We used pooled genomic sequencing to compare male and female genomes between *F. aquilonia*–like and *F. polyctena*–like hybrid lineages and to discover the candidate barrier loci (i.e. genomic regions putatively associated to hybrid male breakdown). We collected the samples used for pooled sequencing from the Långholmen hybrid population (Kulmuni et al. 2010) in the year 2004. The samples were freshly frozen and kept in −20°C and genomic DNA was extracted in the year 2010 from half a body using Qiagen kit. We sequenced altogether four pools of individuals, where each pool consisted of the following number of individuals; 1) 24 *F. aquilonia*–like males, 2) 24 *F. polyctena*–like males, 3) 24 *F. polyctena*–like females and 4) 24 *F. aquilonia*–like females. The sample concentrations were checked with Qubit and pooled into the four pools in equal amounts. Each pool was sequenced with 100bp paired-end sequencing on its own lane in Illumina HiSeq2000 in the Institute for Molecular Medicine Finland (FIMM). This resulted in 46,106,000 to 108,204,481 total number of reads per pool. We quality trimmed reads by removing up to 20 bp that had phred score < 20 using FASTX-Toolkit. Next, we made denovo assemblies of each of the four samples with Soapdenovo trying out different kmer sizes (31, 41, 51, 61, 71) for each assembly. The *F. polyctena*–like male assembly with kmer size of 41 was best in terms of completeness and quality (genome size: 222.6 Mb, 327480 contigs, average contig length: 679 bp, N50 = 1748 bp) and chosen as our reference assembly. We then mapped each sample back to the *F. polyctena*–like male reference assembly after removing contigs of the assembly shorter than 500bp using Bowtie2 v2.0.2. Reads mapped in proper pairs and with a mapping quality superior to 20 were filtered and combined in a single mpileup file using samtools 1.4. Since coverage of the *F. aquilonia*–like male pool was low (mean = 16, s.d. = 38), overlaps between read pairs were kept. For all SNPs (see below) and in each pool, read counts were compared with or without filtering of read pair overlaps using χ^2^ tests. Over all pools, four SNPs displayed significant allele frequency change (*P* < 0.05, for stringency no Bonferroni correction was applied) and were removed from the dataset (Supplementary Figure 5). The mpileup file was then converted in a synchronized file using Popoolation2 [35]. Indels and their 5-bp flanking sequences were masked, and only bi-allelic sites displaying a minimum base quality of 20, coverage between 20 and 60 for each population and a minor allele count of four across all populations were considered. These steps led to the identification of 166,167 SNPs for which *F*_ST_ estimates were computed from read count data using Popoolation2, adjusting for differences in ploidy levels between haploid male and diploid female pools. These estimates were compared with those obtained following a more recently developed approach [36]: both methods provided similar results (ρ = 0.96) and identified a similar number of SNPs differentially fixed in males (Popoolation2: 711; Poolfstat: 719, including all the loci found using Popoolation2). Popoolation2 results were more conservative (i.e, less differentially fixed SNPs) and were used for the rest of the study. For the comparison between poolseq and SNP genotyping data sets, allele counts were imputed from read counts using a maximum-likelihood approach [37], considering the number of chromosomes sampled per pool.

### Gene Annotation, protein-protein interactions and GO term Enrichment Analysis

We defined barrier loci as SNP positions where the male genomes (sampled in 2004) from the *F. aquilonia*–like and *F. polyctena*–like lineages were fixed differently (i.e. *F*_ST_ = 1, see Supplementary Figure 2), which led to identification of 711 SNPs. These fall into to 610 assembled scaffolds. We created an automated pipeline to annotate genomic regions around candidate barrier SNPs, taking advantage of the recent release of the closely related *F. exsecta* genome [17]. First, using our assembly we extracted a sequence of 5,000 bp centered on each of the 711 outlier SNPs. If the contig was smaller than 5,000 bp, the full contig sequence was recovered. Since some SNPs located on the same contig, and sometimes less than 5 kb apart, 639 unique sequences were recovered and were blasted against the *F. exsecta* assembly. For each queried sequence, hits covering less than 60% of its length were filtered out and the best hit was kept per query based on e-value (over all queries, *e*-value < 10^−141^). This allowed to anchor 571 genomic regions on the *F. exsecta* assembly. The coordinates of these regions were extended if needed to reach 5,000 bp, so that sampling effort was equal among genomic regions. *F. exsecta* genes overlapping with best hits were collected using the GenomeIntervals Bioconductor package v1.38. Overall, 590 unique genes overlapped with our 571 genomic regions. Sixty-three of the candidate genomic regions did not contain any annotated gene. The mean number of genes per candidate genomic region was 1.4, and the maximum was six. For each gene, its *F. exsecta* transcript sequence was recovered and blasted against *D. melanogaster* r6.15 proteins after filtering for the longest protein per gene. Alignments below 150 bp and with less than 35% identity were filtered out and the best hit was kept for each query based on e-value (over all queries, *e*-value < 10^−20^). This pipeline recovered 317 *D. melanogaster* genes whose homologs reside in candidate barrier regions. These candidate barrier genes were used for subsequent analyses (e.g., GO and PPI enrichment analyses). For a complete list of scaffolds annotated with their gene names see Supplementary Table 1. GO term enrichment analysis was performed using the Gene Ontology Consortium tool and protein interactions were analyzed using STRING database and *D. melanogaster* homologs for our candidate genes. We performed 1,000 simulations to assess whether the observed PPI enrichment indicated by STRING was significantly different from what would be expected for a random set of SNPs from our dataset. For each simulation, we randomly drew 711 SNPs from the total dataset (166,167 SNPs) and annotated their flanking regions using the same automated pipeline and parameter values as presented above. The number of genes recovered per simulation varied between 267 and 368 (mean = 319). For all simulations and empirical data, PPI enrichment values were computed with STRING v10 using the STRINGdb Bioconductor package [20], setting the score threshold to 800.

We investigated the location of the candidate barrier SNPs blasting to the genome of closely related *F. exsecta*[17] (277 Mb, 14617 scaffolds, N50 = 997654), as our own genomic assembly is highly fragmented.

### Testing for selection by genotyping candidate barrier and random SNPs

To test for selection acting on candidate barrier SNPs we genotyped a set of candidate and random SNPs from samples collected from the study population in 2014. These samples were used to test if significant allele frequency changes occur between larval and adult stages in candidate barrier SNPs but not in random SNPs, as expected if these barrier SNPs mark genomic regions under selection. Genotyping was done at individual level for a total of 196 individuals (Supplementary table 2). Samples were randomly assigned to 96-well plates for genotyping.

Primer design and genotyping were done at LGC genomics using KASP genotyping chemistry. We randomly chose 350 SNPs from the total SNP dataset. Out of these 180 were appropriate for primer design and genotyping. We further chose 183 candidate barrier SNPs out from the 711 SNPs identified previously. Candidate barrier SNPs had *F*_ST_ of 1 between the male genomes but *F*_ST_ varied between females at these loci. We aimed at variable *F*_ST_ estimates between females at the genotyped barrier SNPs and initially chose the 50 SNPs with lowest *F*_ST_, 50 SNPs with highest *F*_ST_ and 100 randomly chosen from the remaining (Figure S1). Of these, 144 SNPs were suitable for genotyping. Therefore, another 106 candidate barrier SNPs were extracted, of which 76 passed the LGC primer design pipeline. In total 220 candidate barrier SNPs were suitable for genotyping and from these we chose 183 SNPs (Supplementary Figure 1). After removal of SNPs or individuals with more than 10% missing data, diploid males from the *F. polyctena*–like lineage and three ambiguous individuals (see Supplementary Figure 3), we were left with 181 individuals (for the *F. polyctena*–like females, early stage: 27, late stage: 32, for the *F. aquilonia*–like individuals, males early stage: 32, late stage: 32, females early stage: 31, late stage: 31) genotyped at 300 SNPs (137 random and 163 putative barrier loci).

Linkage disequilibrium could bias our results if putative barrier SNPs are non-independent or if putative random loci are actually located close to barrier regions. We took advantage of the more contiguous *F. exsecta* genome assembly and pruned the SNP genotyping data based on the location of BLAST hits in the *F. exsecta* genome (see above for BLAST parameters). Since the extent of linkage disequilibrium decay is unknown, we considered three different pruning intervals for SNPs which located on the same *F. exsecta* scaffolds: 50 kb (before pruning: *N*_barrier_ = 163, *N*_random_ = 137; after pruning: *N*_barrier_ = 137, *N*_random_ = 127), 100 kb (*N*_barrier_ = 121, *N*_random_ = 122) and 500kb (*N*_barrier_ = 80, *N*_random_ = 100). We then tested if this had impact on our estimates for *F*_IS_ in females and allele frequency change in males between early and late developmental stages by re-running analysis with the different pruned data sets.

### Statistical analyses

We tested for significant differences in allele counts in *F. aquilonia*–like males between larval and adult stages at candidate barrier SNPs compared to the random SNPs using generalized linear mixed effects model in R [38]. We used locus type (candidate or barrier) and developmental stage (larva or adult) as explanatory variables and locus_ID as a random factor. *F*_IS_ values were estimated per genotyped locus and per group using the HierFstat package [39]. To test if *F*_IS_ estimates were significantly different at candidate barrier loci in females between larval and adult stages we used linear mixed effects model [38] and locus type, developmental stage and group (*F. polyctena* –like or *F. aquilonia*–like) as explanatory variables and locus_ID as a random factor.

## Acknowledgements

We thank Pekka Pamilo and Simon H. Martin for their valuable comments on the manuscript, Petri Kemppainen for discussions and CSC – IT Center for Science, Finland, for computational resources. JK was supported by Human Frontier Science Program Long-term fellowship and Academy of Finland grant nro. 309580. JK and RB designed the study. JK, PN, LP, IS and KD analyzed the data. JK, PN, LP, IS and RB wrote the manuscript. Data will be deposited into Dryad upon acceptance.

