## Supplementary Material for "Evidence for natural selection and barrier leakage in candidate loci underlying speciation in wood ants"

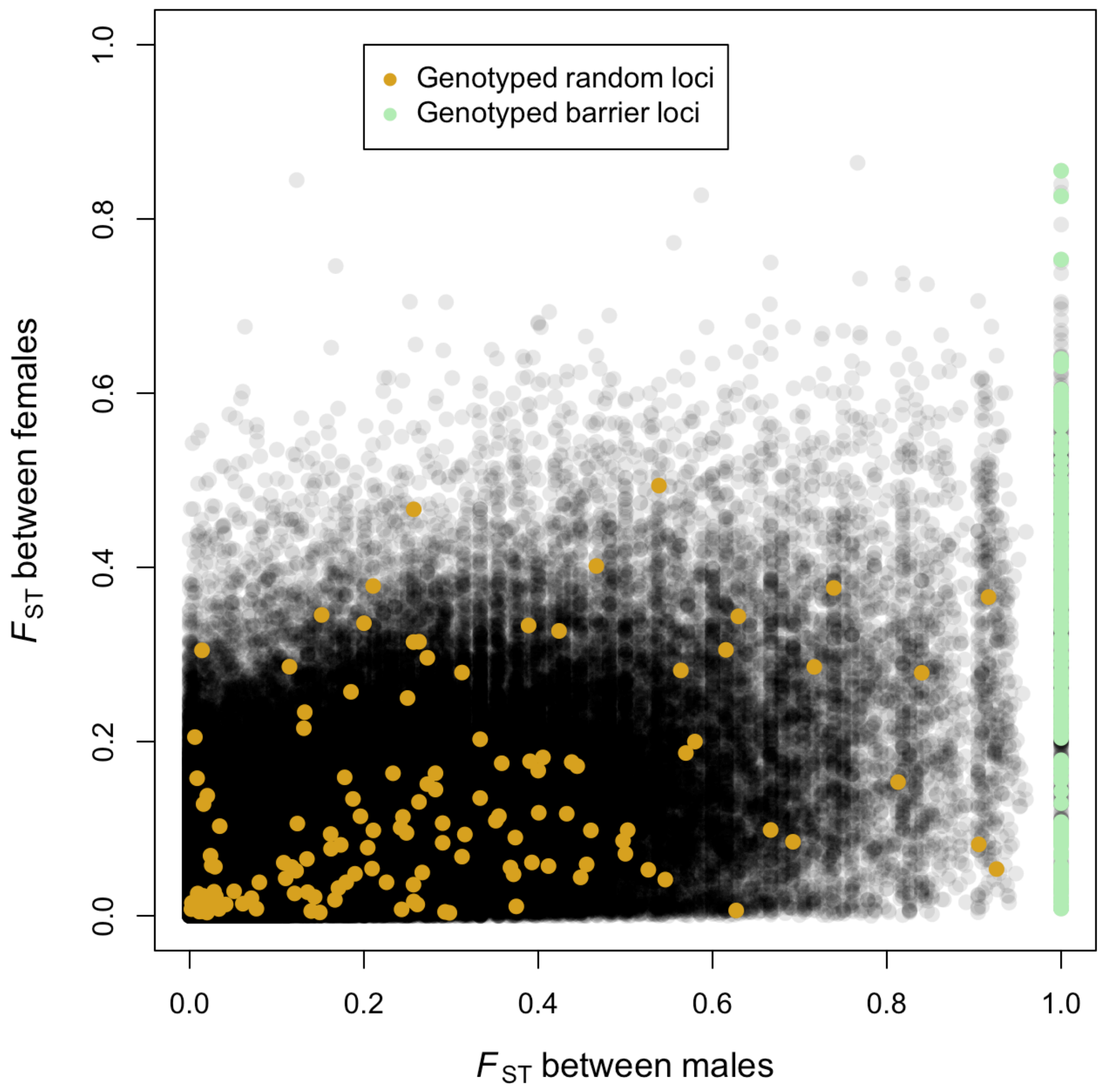


**Supplementary Figure 1.** Genetic differentiation ($F_{ST}$) between the *F. polyctena*-like and *F. aquilonia*-like male genomes (x-axis) versus genetic differentiation between *F. polyctena*-like and *F. aquilonia*-like female genomes (y-axis) for 166,167 SNPs called from pooled sequencing (data collected in 2004). The loci genotyped are colored following; candidate barrier loci are in light green and random loci are in yellow.

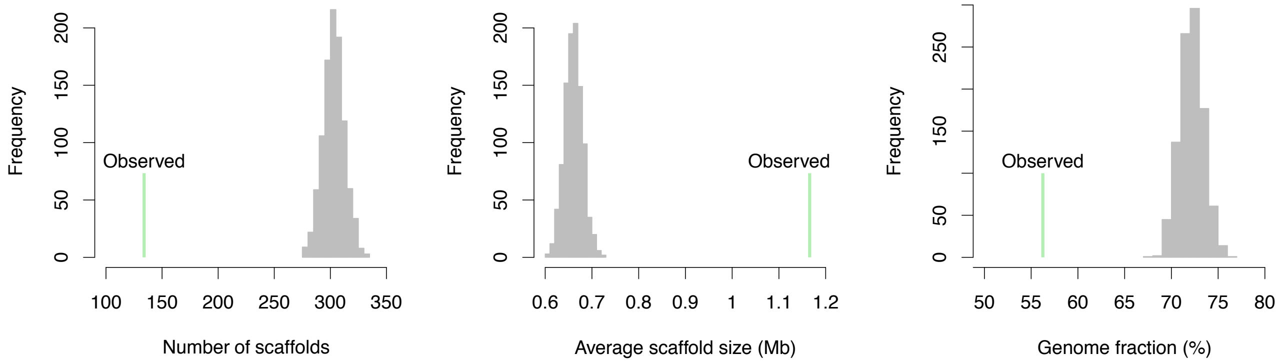


**Supplementary Figure 2.** The candidate barrier loci are more clustered (observed) than randomly drawn polymorphisms in the poolseq data (repeated 1000 times) when mapped to the genome of closely related *F. exsecta*. Left panel; the number of scaffolds the barrier loci locate at, Center panel; average scaffold size, Right panel; cumulative fraction of the genome where candidate barrier loci situate.

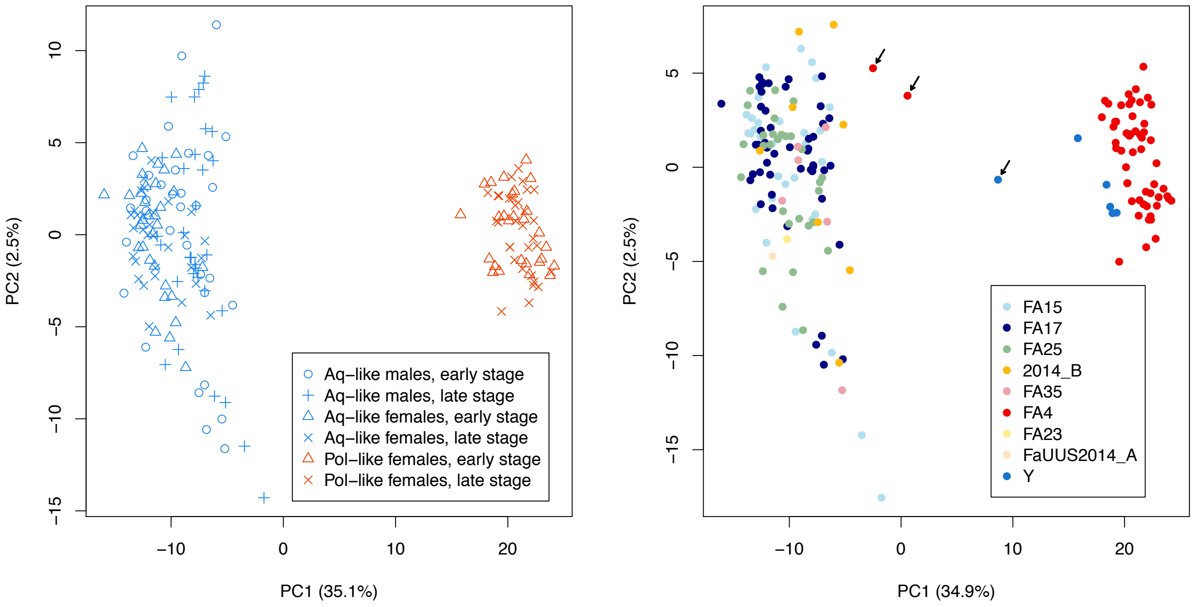


**Supplementary Figure 3.** PCA for 2014 dataset using SNP genotypes with individuals as datapoints. PCA shows no substructure based on nests within *F. aquilonia*-like lineage (left cluster). *F. polyctena*-like (right cluster) individuals come mostly from one nest. a) final data set used in all analyses, b) all individuals colored by nest of origin. Three female larvae (indicated with arrows) found intermediate between *F. aquilonia*-like and *F. polyctena*-like clusters were removed from $F_{IS}$ analysis. The two individuals from nest FA4 (in red) likely represents hybrids between the lineages, while the intermediate individual from nest Y (in blue) had > 30% missing data.

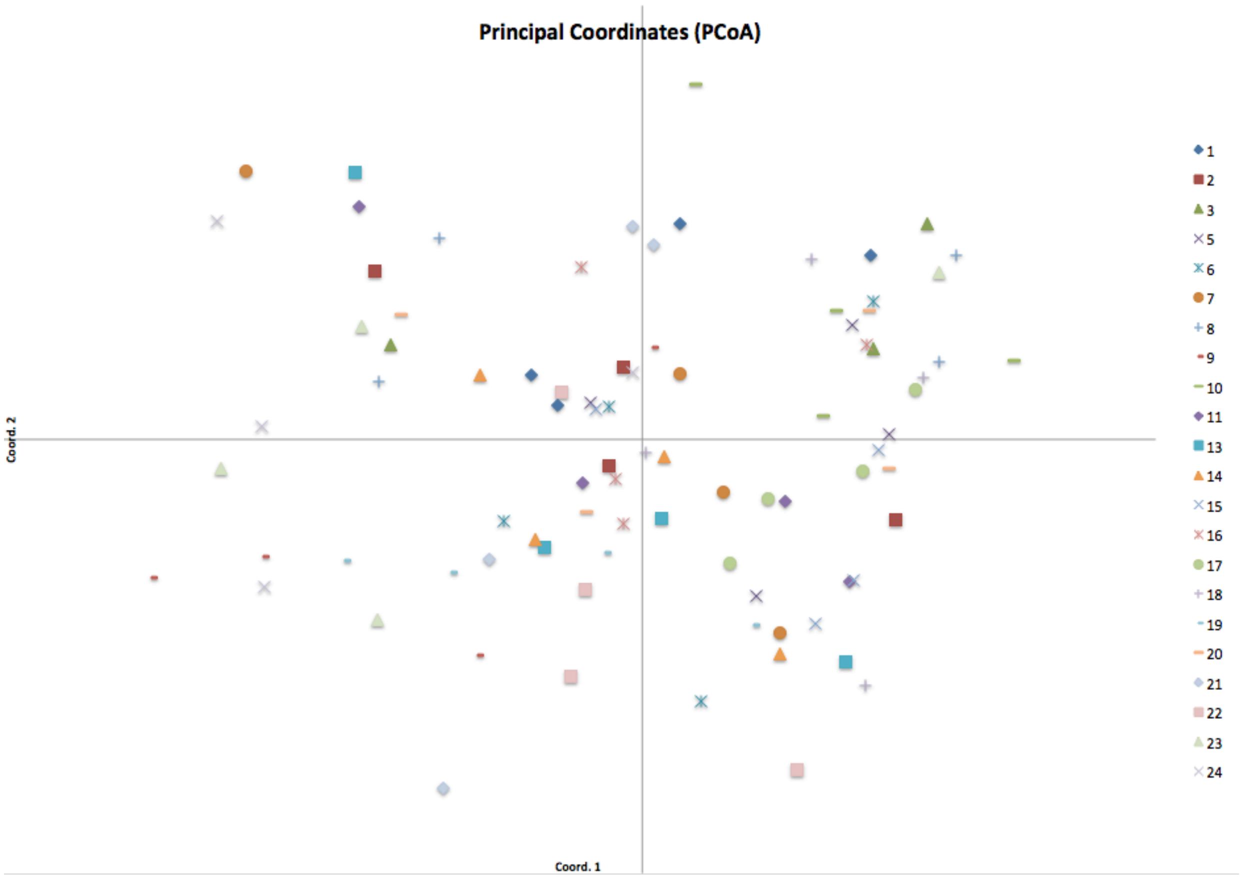


**Supplementary Figure 4.** PcoA shows no substructure in 2005 between individuals coming from different nests of *F. aquilonia*-like lineage based on females (workers) and data from 14 microsatellite markers.

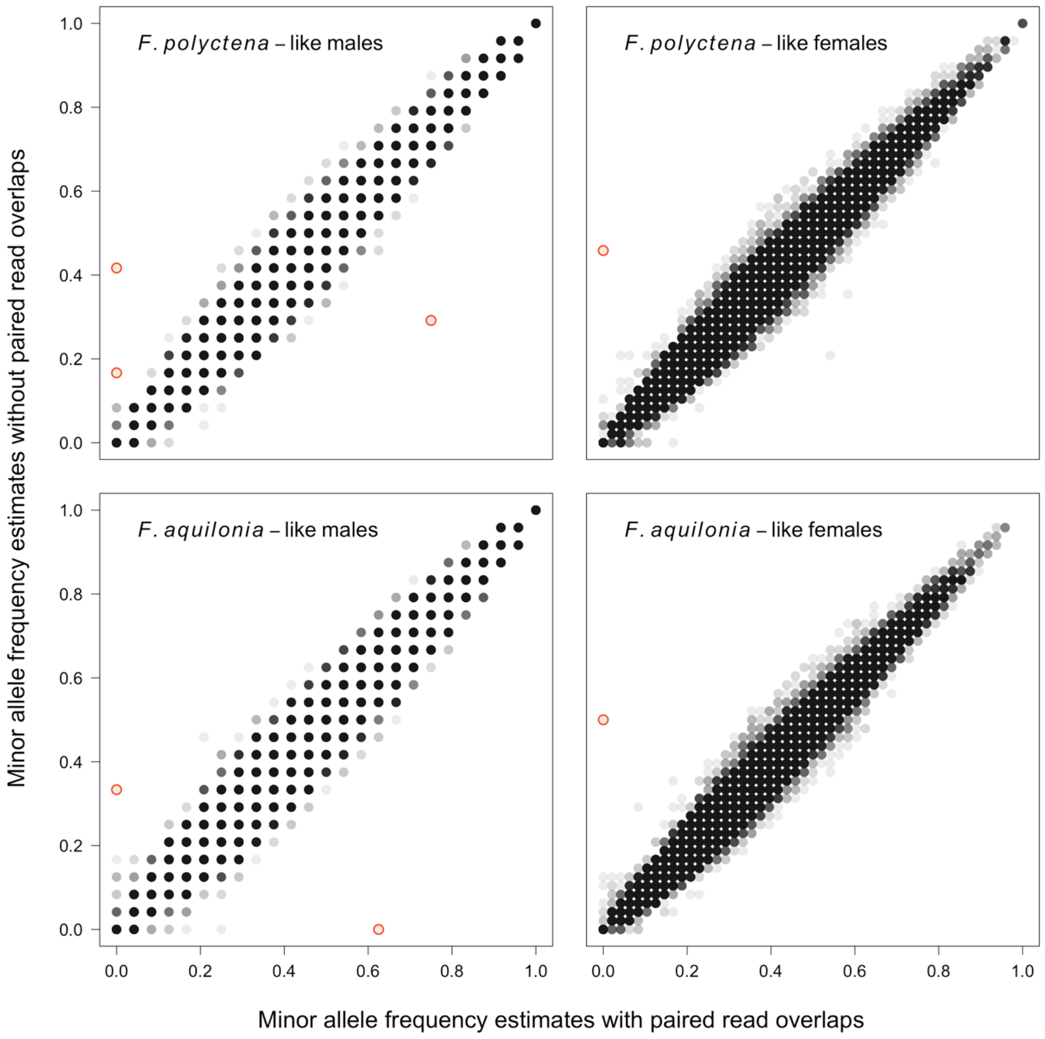


**Supplementary Figure 5.** Impact of read pair overlap filtering on allele frequency estimation in poolseq data. For each pool are plotted the maximum likelihood allele frequency estimates obtained with or without read pair overlap filtering for 166,171 bi-allelic sites displaying a minimum base quality of 20, coverage between 20 and 60 for each population and a minor allele count of four across all pools. SNPs in red display significant differences in read counts between with and without read pair overlap filtering ($P < 0.05$, $\chi^2$ tests) and were removed from the final poolseq SNP dataset.

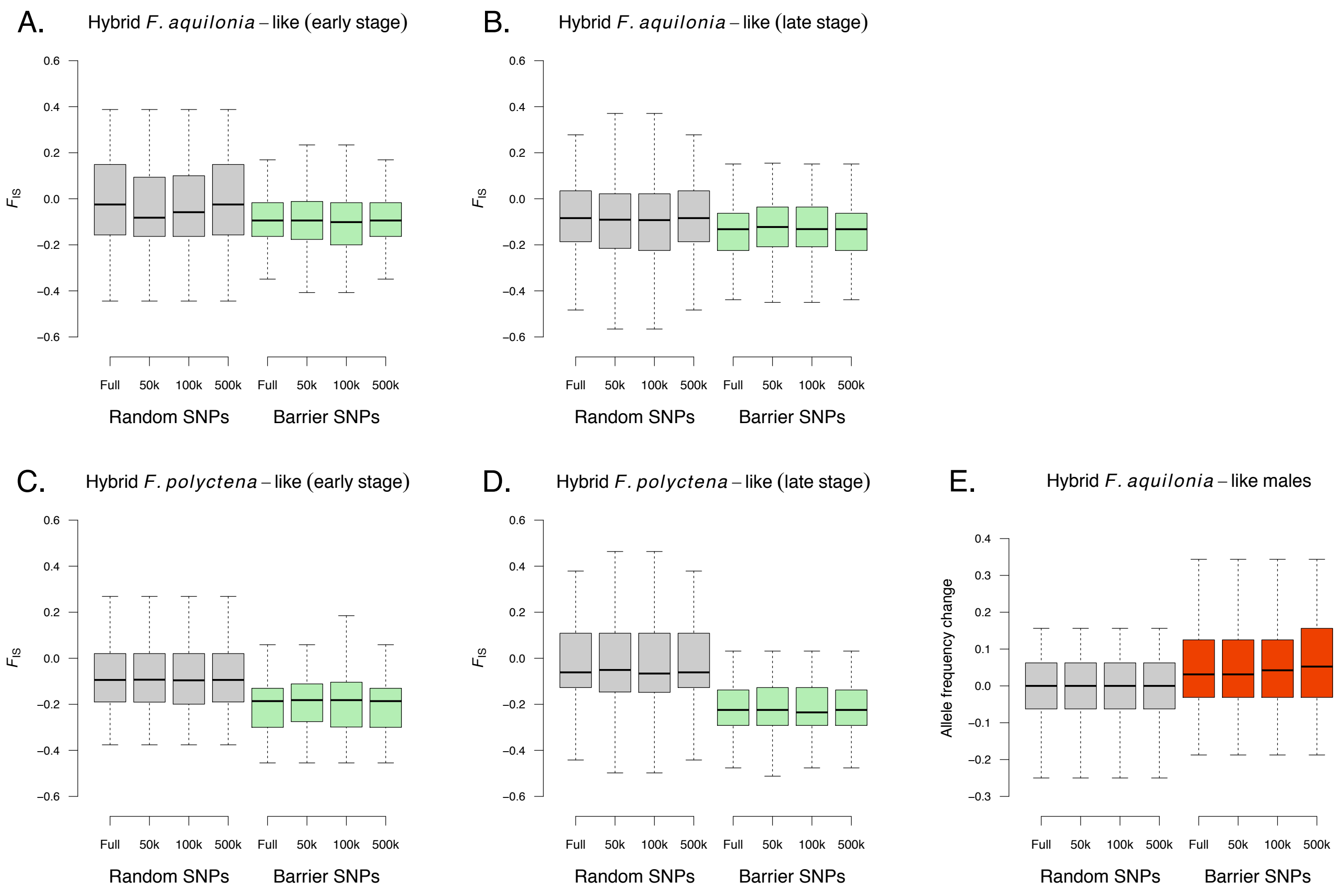


**Supplementary Figure 6.** Effect of SNP pruning on patterns of heterozygosity in females (panels A-D) and allele frequency change in males (panel E) between early and late developmental stages. Full: full data set ($N_{\mathrm{barrier}}$ = 163, $N_{\mathrm{random}}$ = 137), 50k: one SNP is kept every 50 kb ($N_{\mathrm{barrier}}$ = 137, $N_{\mathrm{random}}$ = 127), 100k: one SNP is kept every 100 kb ($N_{\mathrm{barrier}}$ = 121, $N_{\mathrm{random}}$ = 122), 500k: one SNP is kept every 500 kb ($N_{\mathrm{barrier}}$ = 80, $N_{\mathrm{random}}$ = 100). SNP locations were determined after BLAST against the *F. exsecta* genome assembly.

**Supplementary table 1.** Loci annotated in candidate barrier regions. (Attached to the submission as .xls)

**Supplementary table 2.** Samples used for SNP genotyping and testing for selection at candidate barrier loci between early (larval) and late (adult) developmental stages. Samples collected in spring 2014.

| Group | Total number of individuals | Number of individuals per nest |
|---|---|---|
| *F. polyctena*–like males early | 3 | 2 (Nest 4) + 1 (Nest Y) |
| *F. polyctena*–like males late | 5 | 5 (Nest 4) |
| *F. polyctena*–like females early | 27 | 22 (Nest 4) + 5 (Nest Y) |
| *F. polyctena*–like females late | 32 | 32 (Nest 4) |
| *F. aquilonia*–like males early | 32 | 7 (Nest 2014_B) + 6 (Nest 15) + 7 (Nest 17) + 7 (Nest 25) + 5 (Nest 35) |
| *F. aquilonia*–like males late | 32 | 13 (Nest 15) + 18 (Nest 17) + 1 (Nest 25) |
| *F. aquilonia*–like females early | 31 | 1 (Nest 2014_B) + 3 (Nest 15) + 12 (Nest 17) + 1 (Nest 23) + 12 (nest 25) + 1 (Nest 35) + 1 (Nest 2014_A) |
| *F. aquilonia*–like females late | 31 | 10 (Nest 15) + 12 (Nest 17) + 9 (Nest 25) |

**Supplementary table 3.** Allele frequencies at 9 microsatellite loci in the hybrid population in years 1996, 2004, 2008, 2011 and 2014. Data from year 2004 is from ref. Kulmuni et al. 2010 and data from years 2008 and 2011 are from ref. Kulmuni & Pamilo 2014.

| | | workers | workers | adult male | adult gyne | adult male | adult gyne | adult males | adult gynes* | adult males | adult males | adult males | adult gynes in cocoos | adult males* | adult gynes* | adult gynes* |
|---|---|---|---|---|---|---|---|---|---|---|---|---|---|---|---|---|
| | Year | 1996 | 1996 | 2004 | 2004 | 2004 | 2004 | 2008 | 2008 | 2008 | 2011 | 2011 | 2011 | 2014 | 2014 | 2014 |
| **Locus** | **Allele** | **F. aquil. ♀** | **F. polyc. ♀** | **F. aquil. ♂** | **F. aquil. ♀** | **F. polyc. ♂** | **F. polyc. ♀** | **F. aquil. ♂** | **F. aquil. ♀** | **F. polyc. ♂** | **F. aquil. ♂** | **F. polyc. ♂** | **F. polyc. ♀** | **F. aquil. ♂** | **F. aquil. ♀** | **F. polyc. ♀** |
| **FE13** | **N** | 80 | 9 | 62 | 70 | 35 | 25 | 41 | 100 | 41 | 28 | | 34 | 29 | 31 | 32 |
| | **186** | 0,39 | 0,06 | 0,52 | 0,35 | 0,51 | 0,28 | 0,46 | 0,32 | 0,51 | 0,39 | 0,52 | 0,41 | 0,241 | 0,339 | 0,313 |
| | **189** | 0,00 | 0,11 | 0,00 | 0,00 | 0,49 | 0,22 | 0,00 | 0,00 | 0,49 | 0,00 | 0,48 | 0,28 | 0,000 | 0,000 | 0,406 |
| | **198** | 0,66 | **0,83** | 0,48 | 0,65 | **0,00** | **0,50** | 0,54 | 0,68 | **0,00** | 0,61 | **0,00** | **0,31** | 0,759 | 0,661 | **0,281** |
| **FE17** | | | | | | | | | | | | | | | | |
| | **110** | 0,41 | **0,06** | 0,18 | 0,42 | **0,00** | **0,28** | 0,24 | 0,39 | **0,00** | 0,25 | **0,00** | **0,10** | 0,483 | 0,339 | **0,078** |
| | **116** | 0,59 | 0,78 | 0,82 | 0,58 | 0,89 | 0,62 | 0,76 | 0,61 | 0,85 | 0,75 | 0,78 | 0,61 | 0,517 | 0,661 | 0,516 |
| | **118** | 0,00 | 0,17 | 0,00 | 0,00 | 0,11 | 0,10 | 0,00 | 0,00 | 0,15 | 0,00 | 0,22 | 0,29 | 0,000 | 0,000 | 0,406 |
| **FE19** | | | | | | | | | | | | | | 29 | 31 | 32 |
| | **178** | 0,15 | 0,00 | 0,34 | 0,24 | 0,00 | 0,00 | 0,15 | 0,16 | 0,00 | 0,21 | 0,00 | 0,00 | 0,034 | 0,306 | 0,000 |
| | **184** | 0,54 | 0,17 | 0,32 | 0,48 | 0,20 | 0,50 | 0,42 | 0,51 | 0,32 | 0,39 | 0,35 | 0,24 | 0,483 | 0,403 | 0,078 |
| | **186** | 0,17 | 0,61 | 0,32 | 0,11 | 0,80 | 0,38 | 0,15 | 0,21 | 0,68 | 0,36 | 0,65 | 0,71 | 0,207 | 0,145 | 0,750 |
| | **187** | 0,00 | 0,06 | - | - | - | - | - | - | - | - | - | - | - | - | - |
| | **191** | 0,13 | 0,11 | 0,02 | 0,17 | 0,00 | 0,12 | 0,29 | 0,14 | 0,00 | 0,04 | 0,00 | 0,04 | 0,276 | 0,145 | 0,172 |
| | **193** | 0,01 | 0,06 | - | - | - | - | - | - | - | - | | - | - | - | - |
| **FE7** | | | | | | | | | | | | | | | | |
| | **56** | 0,19 | 0,28 | 0,00 | 0,13 | 0,00 | 0,27 | 0,30 | 0,14 | 0,00 | 0,11 | 0,00 | 0,12 | 0,241 | 0,161 | 0,094 |
| | **66** | 0,21 | 0,00 | 0,20 | 0,20 | 0,00 | 0,00 | 0,08 | 0,14 | 0,00 | 0,11 | 0,00 | 0,00 | 0,034 | 0,177 | 0,000 |
| | **68** | 0,58 | **0,06** | 0,80 | 0,67 | **0,00** | **0,23** | 0,63 | 0,73 | **0,00** | 0,78 | **0,00** | **0,06** | 0,724 | 0,661 | **0,109** |
| | **70** | 0,00 | 0,11 | 0,00 | 0,00 | 0,23 | 0,08 | 0,00 | 0,00 | 0,27 | 0,00 | 0,30 | 0,32 | 0,000 | 0,000 | 0,328 |
| | **72** | 0,00 | 0,06 | 0,00 | 0,00 | 0,50 | 0,25 | 0,00 | 0,00 | 0,61 | 0,00 | 0,52 | 0,37 | 0,000 | 0,000 | 0,281 |
| | **89** | 0,00 | 0,06 | 0,00 | 0,00 | 0,26 | 0,17 | 0,00 | 0,00 | 0,12 | 0,00 | 0,17 | 0,13 | 0,000 | 0,000 | 0,188 |
| | **260** | 0,01 | 0,33 | - | - | - | - | - | - | - | - | - | - | - | - | - |
| | **274** | 0,00 | 0,06 | - | - | - | - | - | - | - | - | - | - | - | - | - |
| | **283** | 0,01 | 0,06 | - | - | - | - | - | - | - | - | - | - | - | - | - |
| **FL29** | | | | | | | | | | | | | | | | |
| | **182** | 0,61 | 0,44 | 0,70 | 0,58 | 0,39 | 0,66 | 0,38 | 0,72 | 0,50 | 0,57 | 0,22 | 0,45 | 0,310 | 0,548 | 0,563 |
| | **188** | 0,39 | **0,22** | 0,30 | 0,42 | **0,00** | **0,16** | 0,63 | 0,28 | **0,00** | 0,43 | **0,00** | **0,21** | 0,690 | 0,452 | **0,109** |
| | **190** | 0,00 | 0,33 | 0,00 | 0,00 | 0,61 | 0,18 | 0,00 | 0,00 | 0,50 | 0,00 | 0,78 | 0,34 | 0,000 | 0,000 | 0,328 |
| **FY12** | | | | | | | | | | | | | | 29 | 31 | 32 |
| | **185** | 0,39 | 0,56 | 0,22 | 0,31 | 0,83 | 0,68 | 0,39 | 0,43 | 0,85 | 0,32 | 0,70 | 0,68 | 0,379 | 0,452 | 0,656 |
| | **187** | 0,30 | 0,44 | 0,26 | 0,46 | 0,17 | 0,32 | 0,24 | 0,27 | 0,15 | 0,07 | 0,30 | 0,32 | 0,310 | 0,242 | 0,344 |
| | **189** | 0,30 | 0,00 | 0,52 | 0,24 | 0,00 | 0,00 | 0,37 | 0,30 | 0,00 | 0,61 | 0,00 | 0,00 | 0,310 | 0,306 | 0,000 |
| | **191** | 0,01 | 0,00 | - | - | - | - | - | - | - | | | - | | | |
| **FY13** | | | | | | | | | | | | | | | | |
| | **195** | 0,00 | 0,11 | 0,00 | 0,00 | 0,60 | 0,22 | 0,00 | 0,00 | 0,68 | 0,00 | 0,48 | 0,27 | 0,000 | 0,000 | 0,297 |
| | **197** | 0,17 | **0,56** | 0,03 | 0,11 | 0,00 | 0,22 | 0,20 | 0,16 | 0,00 | 0,11 | 0,00 | 0,06 | 0,379 | 0,161 | **0,016** |
| | **199** | 0,49 | 0,33 | 0,44 | 0,64 | 0,40 | 0,56 | 0,49 | 0,48 | 0,32 | 0,50 | 0,52 | 0,68 | 0,310 | 0,597 | 0,688 |
| | **205** | 0,34 | 0,00 | 0,52 | 0,25 | 0,00 | 0,00 | 0,32 | 0,36 | 0,00 | 0,39 | 0,00 | 0,00 | 0,310 | 0,242 | 0,000 |
| **FY15** | | | | | | | | | | | | | | 29 | 31 | 32 |
| | **221** | 0,34 | 0,06 | 0,40 | 0,42 | 0,18 | 0,08 | 0,30 | 0,35 | 0,23 | 0,52 | 0,17 | 0,39 | 0,241 | 0,355 | 0,359 |
| | **222** | 0,12 | 0,17 | **0,00** | **0,07** | 0,50 | 0,50 | **0,28** | **0,11** | 0,44 | **0,04** | 0,57 | 0,52 | **0,241** | **0,113** | 0,422 |
| | **232** | 0,39 | 0,11 | 0,29 | 0,38 | **0,00** | **0,33** | 0,33 | 0,42 | **0,00** | 0,30 | **0,00** | **0,10** | 0,379 | 0,355 | **0,219** |
| | **234** | 0,14 | 0,00 | 0,31 | 0,13 | 0,00 | 0,00 | 0,10 | 0,12 | 0,00 | 0,15 | 0,00 | 0,00 | 0,138 | 0,177 | 0,000 |
| | **236** | 0,00 | 0,06 | - | - | - | - | - | - | - | - | - | - | - | - | - |
| | **240** | 0,00 | 0,06 | 0,00 | 0,00 | 0,32 | 0,08 | 0,00 | 0,00 | 0,33 | 0,00 | 0,26 | 0,00 | - | - | - |
| | **244** | 0,01 | 0,56 | - | - | - | - | - | - | - | - | - | - | - | - | - |
| **FY3** | | | | | | | | | | | | | | | | |
| | **185** | 0,61 | 0,44 | 0,54 | 0,57 | 0,80 | 0,58 | 0,71 | 0,62 | 0,90 | 0,75 | 0,87 | 0,52 | 0,483 | 0,629 | 0,500 |
| | **188** | 0,17 | 0,00 | 0,46 | 0,25 | 0,00 | 0,00 | 0,20 | 0,19 | 0,00 | 0,18 | 0,00 | 0,00 | 0,069 | 0,242 | 0,000 |
| | **190** | 0,22 | 0,56 | **0,00** | **0,18** | 0,20 | 0,42 | **0,10** | **0,19** | 0,10 | **0,07** | 0,13 | 0,49 | **0,448** | **0,129** | 0,500 |
| | **194** | 0,01 | 0,00 | - | - | - | - | - | - | - | - | | - | | | |

All individuals indicated as males and gynes are individuals hatched during that season. Workers were collected from year 1996.
*adult individuals and adults inside cocoos ready to hatch
Orange background indicates alleles diagnostic to F. polyctena and blue background alleles diagnostic to F. aquilonia
Alleles bolded orange are F. polyctena alleles introgressed to F. aquilonia and alleles bolded blue are F. aquilonia alleles introgressed to F. polyctena.
